# SLC7A11 is a superior determinant of APR-246 (Eprenetapopt) response than *TP53* mutation status

**DOI:** 10.1101/2020.11.29.398875

**Authors:** Kenji M. Fujihara, Mariana Corrales-Benitez, Carlos S. Cabalag, Bonnie Z. Zhang, Hyun S. Ko, David S. Liu, Kaylene Simpson, Ygal Haupt, Sue Haupt, Wayne A. Phillips, Nicholas J. Clemons

## Abstract

**Purpose:** APR-246 (Eprenetapopt) is in clinical development with a focus on haematological malignancies and is marketed as a mutant-p53 reactivation therapy. Currently, the detection of at least one *TP53* mutation is an inclusion criterion for patient selection into most clinical trials. Preliminary results from our phase Ib/II clinical trial investigating APR-246 combined with combination chemotherapy (cisplatin and 5-Fluorouracil) in metastatic oesophageal cancer, together with previous pre-clinical studies, indicate that *TP53* mutation status alone may not be a sufficient biomarker for response to APR-246. This study aimed to identify a robust biomarker for response to APR-246.

**Methods:** Correlation analysis of the PRIMA-1 activity (lead compound to APR-246) with mutational status, gene expression, protein expression and metabolite abundance across over 800 cancer cell lines was performed. Functional validation and a boutique siRNA screen of over 750 redox-related genes were also conducted.

**Results:** *TP53* mutation status was not predictive of response to APR-246. The expression of SLC7A11, the cystine/glutamate transporter, was identified as a superior determinant of response to APR-246. Genetic regulators of SLC7A11, including ATF4, MDM2, wild-type p53 and c-Myc were confirmed to also regulate cancer cell sensitivity to APR-246.

**Conclusions:** SLC7A11 expression is the major determinant of sensitivity to APR-246 and should be utilised as a predictive biomarker in future clinical investigation of APR-246.

## INTRODUCTION

Detailed understanding of drug mechanism-of-action is vital for assessing on-target efficacy *in vivo*, to facilitate the translation of efficacious, novel anti-cancer therapies into the clinic. Strong predictive biomarker identification can then follow, allowing for rational patient selection for clinical trials, which in turn dictates appropriate application in the clinic. APR-246 is proposed to be a mutant-p53 (mut-p53) reactivator and is currently under clinical investigation in *TP53*-mutated myelodysplastic syndrome (MDS) and acute myeloid leukaemia (AML)^1^. The mechanism-of-action of APR-246 remains contentious. APR-246, like its lead compound PRIMA-1, is a pro-drug that is converted to the active compound methylene quinuclidinone (MQ) that conjugates to thiols through Michael Addition^2^. The predominant reported mechanism-of-action involves reactivation of wild-type p53 (wt-p53) activity through covalent modification of cysteine residues in the core domain of mut-p53 protein^2,3^. However, more recent work links the anti-neoplastic capacity of APR-246 to targeting antioxidant pathways, causing glutathione (GSH) depletion and direct conjugation and inhibition of enzymes involved in resistance to oxidative stress^4,5^.

Currently, the vast majority of APR-246 clinical trials include the detection of at least one *TP53* mutation as inclusion criteria. The variable mut-p53 dependence for efficacy of APR-246, observed in some studies^6,7^, has been rationalised as cancer-type specific, however no unequivocal predictors of response to APR-246 in a pan-cancer setting are known. More recently, *ARID1A* mutation was also shown to drive sensitivity to APR-246 and other GSH depleting agents^8,9^. Previous cellular profiling efforts combining small-molecule sensitivity data with cancer cell line omics datasets, in order to identify unique and robust biomarkers for small molecule activity have been successful^10^. Identification of a strong predictive biomarker for this first-in-class compound APR-246, which is indicative of its true mechanism-of-action, will help to ensure its continued clinical development.

Here, using a pan-cancer, cellular-feature, correlative approach to incorporate transcriptomic, metabolomics and proteomic data, we identified the cystine-glutamate antiporter, SLC7A11, to be a robust biomarker for APR-246. Remarkably, the predictive capacity of SLC7A11 proved to be superior than *TP53* mutation status, highlighting its value as a novel prognostic indicator for patient stratification of APR-246 administration.

## RESULTS

### Clinical investigation of APR-246 in advanced and metastatic oesophageal and gastro-oesophageal junction cancer

The translation of our laboratory’s previous findings, demonstrating efficacy of APR-246 in combination with chemotherapy in pre-clinical models of oesophageal cancer^11^, was launched as an investigator-led phase Ib/II clinical trial to investigate the safety and efficacy of APR-246 in combination with cisplatin and 5-Fluorouracil (5-FU), in advanced and metastatic oesophageal and gastro-oesophageal junction cancer^12^. Five patients completed enrolment for the phase I trial, before its suspension. All patients received Level 1 dose regimen. Given the difficulties inherent in accessing patient tumour materials directly in the metastatic setting, analysis of efficacy and biomarker investigations were undertaken using radiographic imaging and circulating tumour DNA (ctDNA) (**Figure 1A**). Somatic mutations were detected in four out of five patients by ctDNA at baseline, using a targeted amplicon panel of nine genes that are commonly mutated in oesophageal cancer^13^. Of importance to APR-246, *TP53* mutations were detected in 3 patients, whilst *ARID1A* mutations were identified in two patients. Intriguingly, compared to those patients with detectable *TP53* and/or *ARID1A* mutations who progressed within the first 2 cycles, patient PMC002 showed stable disease over 5 trial cycles, yet did not have any detectable *TP53* or *ARID1A* mutations by ctDNA analysis. PMC002’s tumour burden as determined by serial CT scans of target metastatic lesions was stable – exemplified by their posterior lower paraoesophageal node (**Figure 1B,C)**. Meanwhile, another patient (SUN002), with both detected *TP53* and *ARID1A* mutations, progressed on trial after only two cycles, as evident by both a rise in ctDNA and in their CT scans (**Figure 1D,E**). Although the number of patients is small, these clinical results emphasise the need to establish a sensitive and robust biomarker for APR-246 response.

**Figure 1.**
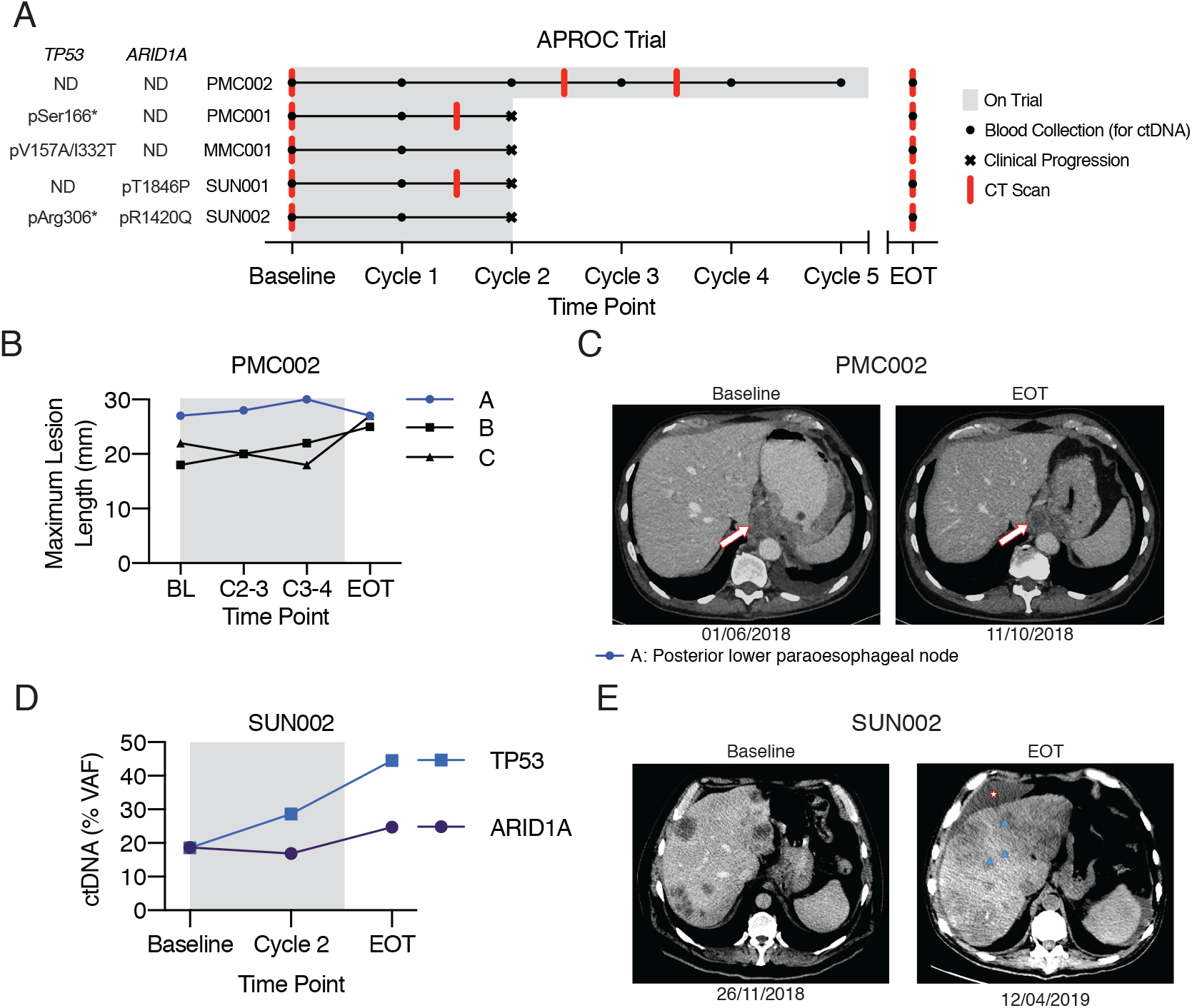
Clinical investigation of APR-246 in advanced and metastatic oesophageal and gastrooesophageal junction cancer. (A) Schematic representing the patients recruited into the APR-246 in Oesophageal Cancer (APROC) Phase Ib/II trial indicating number of cycles of APR-246 received in combination with chemotherapy (5-FU and Cisplatin), when CT scans and blood was taken for circulating tumour DNA (ctDNA) to track progression, and indicates the presence of *TP53* and *ARID1A* mutations as detected by ctDNA targeted amplicon sequencing (TAMseq). (B) Maximum dimension of patient target metastatic lesions (by RECIST 1.1) measured by CT scan timepoints and (D) representative CT scan images at indicated timepoints in patient PMC002. (A: Posterior lower paraoesphageal node, B: left lateral supraclavicular node, C: Right level V lower cervical node). (D) Levels of ctDNA (variant allele frequency, VAF) and (E) CT scans at indicated timepoints in patient SUN002. Blue triangles indicate new metastatic lesions, star indicates liver ascites. Data in (B) is represented as mean of two technical replicates.

### Performance of APR-246 in the Cancer Therapeutics Response Portal v2

In order to identify a robust biomarker for response to APR-246, we interrogated the pancancer therapeutic response and cancer cell line omics datasets from the Broad Institute’s DepMap portal (www.depmap.org). Both APR-246 and PRIMA-1 were included in the Broad Institute’s Cancer Target Response Portal version 2 (CTRPv2) small-molecule sensitivity profiling effort^14,15^. However, only PRIMA-1 was profiled at a sufficient concentration to ensure a lethal dose was achieved across a majority of cancer cell lines (**Figure 2A).** As a result, PRIMA-1 and APR-246 showed only a mild similarity in their sensitivity profiles based on area-under-the-curve (AUC) analysis, however single dose effects, at comparable concentrations, highlighted remarkable concordance between sensitivity to APR-246 and PRIMA-1 (**Figure 2B**). Therefore, PRIMA-1 sensitivity profile was utilised for further analyses.

**Figure 2.**
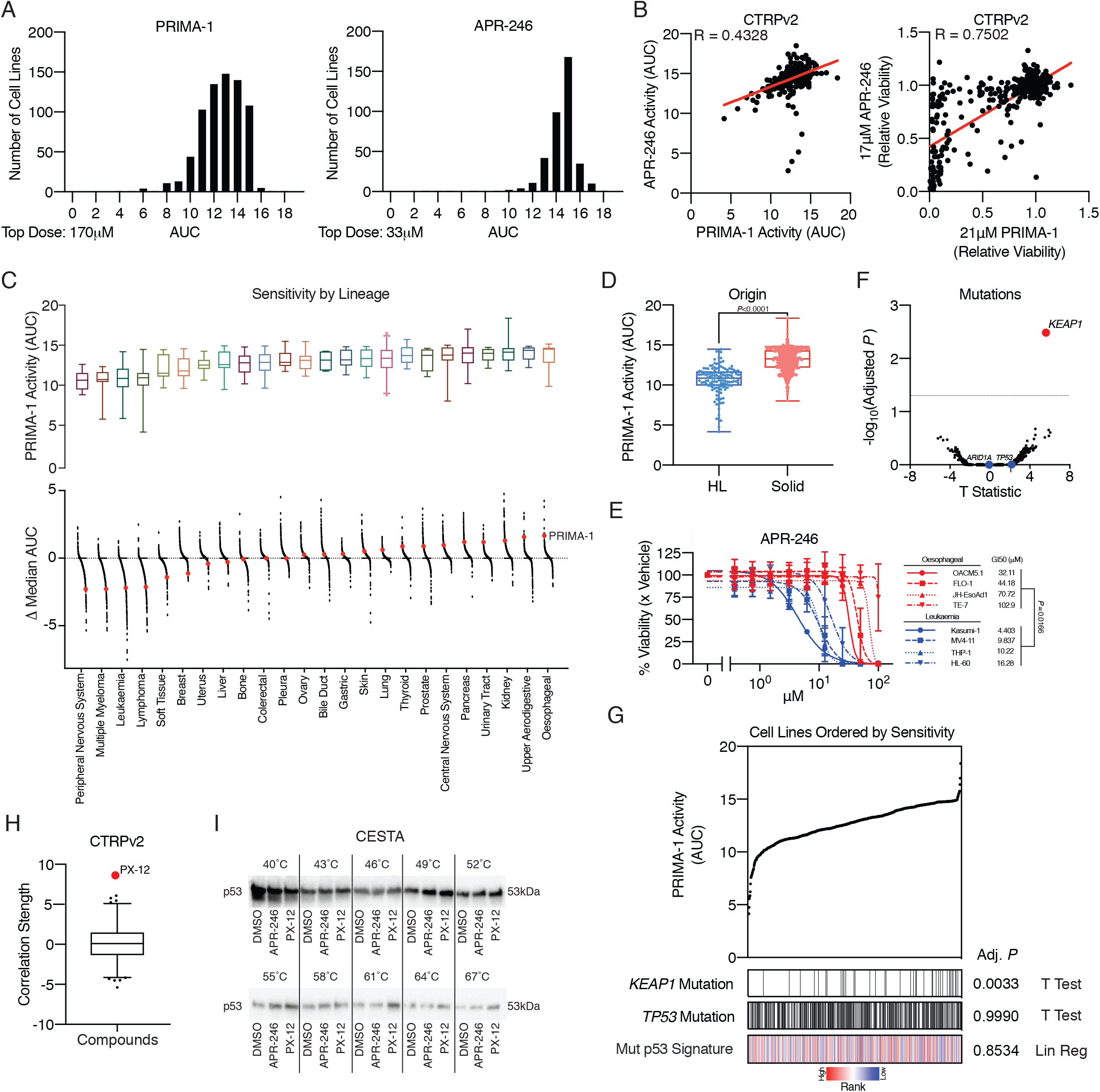
Performance of APR-246 and PRIMA-1 in the Cancer Therapeutics Response Portal v2. (A) Histograms of PRIMA-1 and APR-246 activity in cancer cell lines as measured by area-under-the-curve (AUC). (B) Scatterplots of PRIMA-1 and APR-246 AUC (left) and relative viability at comparable doses (right). (C) Box-and-whisker plots (1^st^-99^th^ percentile) of lineage specific sensitivity to PRIMA-1 ordered by median sensitivity (top), and relative activity of individual compounds by lineage – indicating PRIMA-1 in red (bottom). Points represent the 481 compounds included in the CTPRv2. Points above the dotted line indicate preferentially inactive compounds in a specific lineage, whilst points below the dotted line are preferentially active. (D) Boxplots (min-max) comparing sensitivity to PRIMA-1 in haematopoietic and lymphoma (HL) and solid tumour derived cell lines. (E) Cell viability following 72 h exposure with APR-246 at indicated doses in leukaemia (blue) and oesophageal (red) cancer cell lines. (F) Volcano plot comparing the effect of mutation status on PRIMA-1 sensitivity across all cancer cell lines. Dotted line indicates Benjamin-Hochberg cut off for false discovery. *KEAP1* mutation (red) is significantly associated with resistance to PRIMA-1, *TP53* and *ARID1A* mutation are not significantly association with sensitivity or resistance to PRIMA-1. (G) PRIMA-1 activity and top correlated cancer cell line mutation status (indicated by black lines for mutations), *KEAP1*, as well as *TP53* mutation and mut-p53 transcriptional signature (see methods for details). (H) Box-and-whisker plot (1^st^-99^th^ percentile) of Fischer’s transformed z-scored Pearson correlation strength of PRIMA-1 AUC and the other 480 CTRPv2 compounds, indicating PX-12 as the most significant outlier in red. (I) Cellular thermal shift assay (CETSA) performed in H1299 cells overexpressing mutant-p53 (R273H). Cells were treated for 2 h with DMSO, pre-heated 50 μM APR-246 or 25 μM PX-12 before harvest, freeze-thaw and subject to increasing temperatures. (B, G – Mut p53 signature) Pearson’s correlation, two-tailed unpaired t-test (D, E, F, G). Error bars = SEM. (E) *N*=2-3, (I) *N*=3, representative blot. *See also Supplementary Fig. 1 and 2*.

Cancer cell line (CCL) response data to PRIMA-1 across 24 tissue lineages highlighted higher sensitivity amongst haematopoietic and lymphoid malignancies (multiple myeloma, lymphoma and leukaemia) compared to solid malignancies (**Figure 2C,D**). In keeping with previous studies^15,16^, the 481 compounds in CTRPv2 were generally more efficacious in haematopoietic and lymphoid (HL) cancer cell lines compared with those from solid cancers (**Figure 2C-bottom**). We also confirmed that APR-246 has increased efficacy in representative leukaemia cell lines compared to oesophageal CCLs (**Figure 2E**). Of note, oesophageal CCLs were the most resistant to PRIMA-1 in this analysis, meanwhile peripheral nervous system (PNS) CCLs were the most sensitive.

Next, analysis of the binary mutational association with PRIMA-1 sensitivity across all CCLs was performed (**Figure 2F**). Strikingly, *TP53* or *ARID1A* mutation status were not predictive of response to PRIMA-1, whilst *KEAP1* mutation was associated with resistance **(Figure 2F, G)**. KEAP1 is an E3 ubiquitin-ligase known to control the expression of the master transcriptional regulator of anti-oxidant response, NRF2 (encoded by *NFE2L2*)^17,18^. Importantly, *TP53* mutation status alone is not predictive of favourable response across solid, HL, lung or breast cancer cell lineages, if anything mutant *TP53* is predictive of decreased sensitivity in a pan-cancer analysis and in HL lineages (**Figure S1A**). Additionally, use of a recently published *Mut p53 signature*^19^ based on cancer cell gene expression profiles had no correlation with PRIMA-1 activity (**Figure 2G**). Finally, whilst we previously demonstrated that mut-p53 protein accumulation correlated with increased sensitivity to APR-246 in oesophageal CCLs^4,11^, no comparable correlation was evident in a pan-cancer CCL analysis, based on p53 protein expression as determined by quantitative proteomics (**Figure S 1B**)^20^. This suggests a relationship in oesophageal lines, but led us to look for a universal marker of APR-246 response.

Intriguingly, correlation of PRIMA-1 activity to the other 480 compounds in the CTRPv2, revealed that PX-12, a thioredoxin inhibitor^21^, shares a similar activity profile with PRIMA-1 among CCLs (**Figure 2H, Figure S 2A**). Interestingly, whilst overexpression of mut-p53 (R273H) led to an increase in sensitivity to APR-246 in p53-null H1299, there was no change in sensitivity to PX-12 (**Figure S 2B**). Given that APR-246 has been previously demonstrated to thermostabilise mut-p53 through direct MQ conjugation^3^, we accessed the effect of APR-246 and PX-12 incubation on mut-p53 thermostability in H1299 cells through cellular thermal shift assay^22^. Surprisingly, PX-12 maintained mut-p53 thermostability to a greater extent than APR-246 at comparable efficacious doses (**Figure 2I, Figure S 2C**). Together, these data confirm *TP53* mutation status alone is insufficient to determine sensitivity to APR-246 across multiple lineages of CCLs.

### SLC7A11 is the major determinant of APR-246 sensitivity

To look beyond association with *TP53* mutation status, we performed a range of analyses correlating CCL response to PRIMA-1 with cell line omics features, including gene transcript levels, protein expression and metabolite abundance^20,23,24^. To avoid potential confounding effects of increased sensitivity to PRIMA-1 in HL lineages, solid and HL CCL lineages were separated for these analyses. Strikingly, SLC7A11 mRNA and protein and reduced glutathione (GSH) were the strongest predictive biomarkers for response to PRIMA-1 in solid CCLs (**Figure 3A**). This was consistent across analyses of HL, lung and breast lineages (**Figure S 3A**). SLC7A11, expression of which is regulated predominantly by NRF2^25^, is the key functional subunit of the system x_c_^-^ antiporter, which imports extracellular cystine in exchange for intracellular glutamate^26^. The import of cystine through system x_c_^-^ provides the predominant source of intracellular cysteine, which is the rate limiting substrate required for *de novo* glutathione synthesis^27^. Further, mRNA and protein expression of SLC7A11 is consistently lower in HL compared to solid CCLs (**Figure 3B**), likely driven by lower levels of NRF2 mRNA (**Figure S 3B**) – consistent with the finding that *KEAP1* mutation is predictive of resistance to PRIMA-1 in CCLs (**Figure 2F**). Reliably, significant correlations exist between *SLC7A11* expression and PRIMA-1 activity in both solid and HL CCLs (**Figure 3C**). Validating these findings, APR-246 resistance is induced by the ectopic overexpression of SLC7A11 in H1299 p53^Null^ *KEAP1* wild-type cells (**Figure 3D)**. In contrast, acute knockdown of SLC7A11 prior to APR-246 treatment, increased sensitivity to APR-246. Altogether, these results illustrate that SLC7A11 is the major determinant of APR-246 sensitivity across multiple cancer lineages with varying *TP53* mutational status.

**Figure 3.**
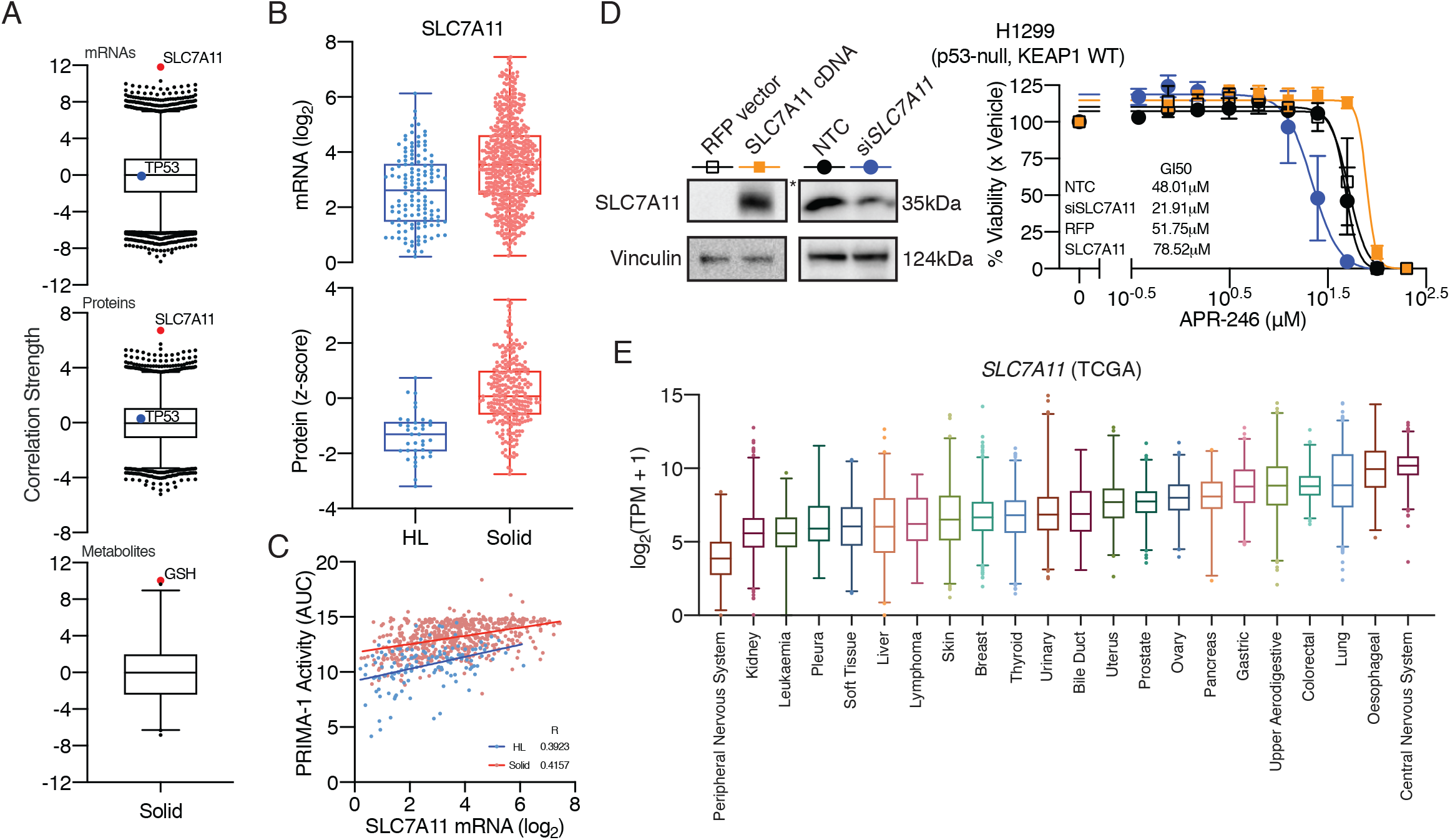
SLC7A11 expression is the major determinant of APR-246 sensitivity. (A) Box-and-whisker plots (1^st^-99^th^ percentile) of Fischer’s transformed z-scored Pearson correlation strengths of PRIMA-1 AUC and gene transcripts, proteins and metabolites in solid cancer. Red dots indicate SLC7A11 mRNA and protein, and glutathione (GSH) correlation strengths. Blue dots indicate TP53 mRNA and protein correlations. (B) Boxplots (min-max) comparing SLC7A11 mRNA and protein levels in HL and solid tumour derived cell lines. (C) Scatterplot correlating the expression of *SLC7A11* with PRIMA-1 AUC in HL (blue) and solid (red) cancer cell lines. (D) SLC7A11 and red fluorescent protein (RFP) stable overexpression and transient *SLC7A11* knockdown in H1299 cells confirmed by western blot (left). * indicates short exposure time. APR-246 was applied 24 h post transfection of *SLC7A11* and nontargeted control (NTC) siRNA or plating of overexpression cells with viability measured at 72 h post APR-246 (right). (E) Box-and-whisker plots (1^st^-99^th^ percentile) of normalised expression of *SLC7A11* across 23 tumour types by lineage in publicly available TCGA datasets. (C) Pearson’s correlation. *See also Supplementary Fig. 3*.

Extending the clinical validity of these findings, we analysed the expression levels of *SLC7A11* across all tumour lineages in The Cancer Genomic Atlas (TCGA) (**Figure 3E**) and found that the SLC7A11 expression across the majority of tumour lineages was generally consistent with the cell line lineage sensitivity to PRIMA-1 (**Figure 2A**). For example, leukaemia samples express low levels of *SLC7A11*, whilst oesophageal tumours have high expression – corresponding with the sensitivity of these lineages to APR-246.

### siRNA screen reveals modulators of SLC7A11 as key genetic regulators of APR-246 sensitivity

To complement our biomarker investigation, we performed a boutique siRNA synergy screen with a library targeting 753 genes in redox metabolism and/or NRF2 target genes (**Figure 4A**). In order to account for mut-p53-dependent and independent effects of APR-246 and redox perturbation synergy, parallel screens were performed in isogenic p53-null and mut-p53 overexpression cancer cells. To this point, there was significant concordance between the two isogenic cell lines across two independent siRNA screen runs (**Figure 4B**). Consistently, *SLC7A11*, as well as *SLC3A2* (another component of system x_c_^-^), were amongst the top gene knockdowns synergistic with APR-246 in both cell lines (**Figure 4B)**. Interestingly, *MDM2* and *ATF4* gene ablation were also synergistic with APR-246, independent of mut-p53 expression (**Figure 4B,C**). MDM2 is the major negative regulator of p53 protein expression – acting as an E3 ligase that targets p53 for proteasome-mediated degradation^28^, however it also has p53-independent activities. Previous efforts investigating p53-independent functions of MDM2 revealed that MDM2, together with ATF4, directly regulate cellular redox status through transcriptional regulation of *SLC7A11*^29^. In keeping with this model, siRNA ablation of *ATF4* or *MDM2* decreased the expression of *SLC7A11* (**Figure 4D**), likely explaining the synergy with APR-246. Importantly, these findings, further demonstrate the importance of SLC7A11 in determining APR-246 sensitivity and highlight that multiple mechanisms can modulate SLC7A11 in cancer cells.

**Figure 4.**
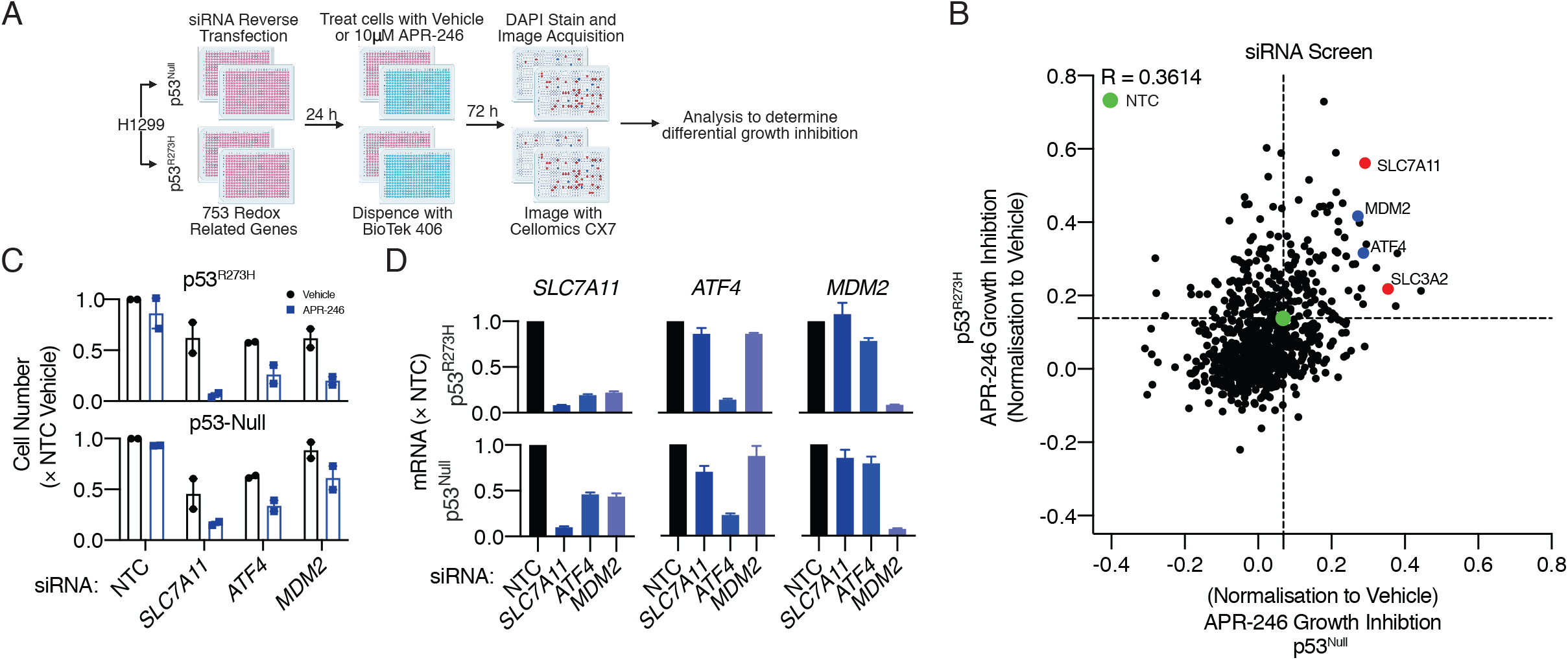
siRNA screen reveals modulators of SLC7A11 expression as key genetic regulators of APR-246 sensitivity. (A) Schematic representation of the boutique siRNA screen performed in H1299 parental cells (p53^Null^) and cells overexpressing mutant-p53 hotspot R273H (see methods for details). (B) Scatterplot indicating APR-246-mediated growth inhibition following siRNA transfection. Dotted line indicates APR-246 (10 μM) growth inhibition on non-targeting control (NTC). (C) Relative cell number changes following 24 h siRNA-mediated knockdown of *SLC7A11, ATF4* and *MDM2* and 72 h treatment with 10 μM APR-246 from the two independent siRNA screen runs. (D) Changes in *SLC7A11, ATF4* and *MDM2* expression following 24 h knockdown. Error bars = SEM. (B, C) *N*=2. (D) *N*=3.

### Wild-type p53 regulation of SLC7A11 is indirect and dictates sensitivity to APR-246

Given that the canonical regulator of p53, MDM2, regulates SLC7A11, we next focused on examining the relationship between wt-p53 and SLC7A11 in relation to APR-246 sensitivity. Using p53-null H1299 cells engineered with a ponasterone-A (Pon-A) inducible wt-p53 construct, we characterised the effects of p53 on SLC7A11 expression. Consistent with previous studies^30,31^, ectopic overexpression of wt-p53 resulted in decreased SLC7A11 mRNA and protein levels (**Figure 5A, Figure S 4A**). Functionally, this resulted in decreased total glutathione levels and increased sensitivity to APR-246 (**Figure 5B, C**). Interestingly, whilst no effect was observed on ATF4 protein levels, wt-p53 overexpression led to stabilisation of NRF2 protein and upregulation of other NRF2 target genes, *HMOX1* and *KEAP1* (**Figure S 4A,B**).

**Figure 5.**
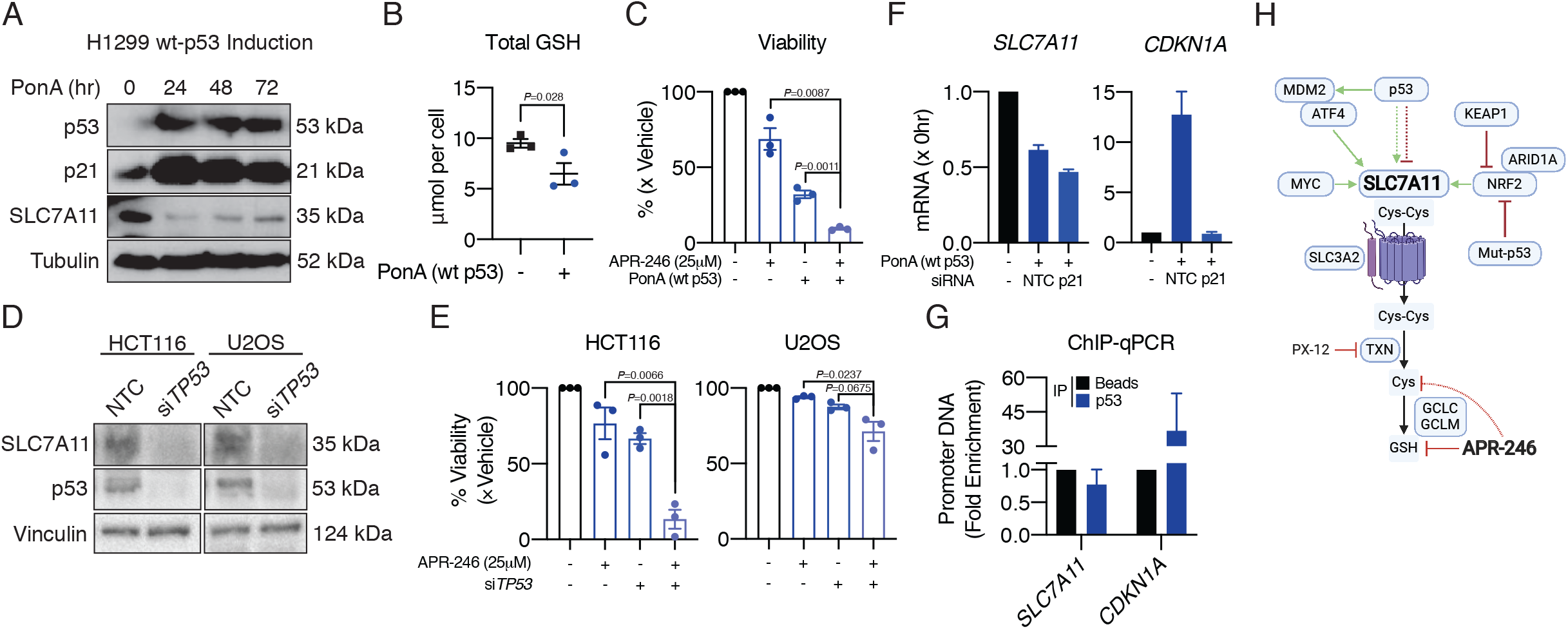
Wild type p53 indirectly regulates SLC7A11 expression and alters sensitivity to APR-246. (A) Western blot of p53, SLC7A11 and p21 protein levels in H1299 cells with Ponasterone A (PonA) inducible wild type p53 (wt-p53) following treatment with 2.5 μg/mL PonA for indicated timepoints. (B) Total GSH levels following 48 h PonA treatment. (C) Viability following 48 h pre-treatment with PonA and subsequent treatment with 25 μM APR-246 or vehicle. (D) p53 and SLC7A11 protein levels following 48 h knockdown with *TP53* siRNA or NTC in wt-p53 HCT116 and U2OS cancer cell lines. (E) Viability following 24 h *TP53* knockdown and subsequent treatment with 25 μM APR-246 or vehicle in wt-p53 HCT116 and U2OS cancer cell lines. (F) *SLC7A11* and *CDNK1A* (p21) mRNA levels following 48 h knockdown with *CKDN1A* or NTC siRNA and co-induction of wt-p53 with PonA in H1299 cells. (G) Chromatin immunoprecipitation (ChIP) of the *SLC7A11* and *CDKN1A* promoters using p53 antibody or agarose control beads following 48 h treatment in H1299 wt-p53 cells. (H) Schematic representation summarising the regulation of SLC7A11 in cancer cells and downstream control of sensitivity to APR-246. Dotted arrows indicate indirect regulation or targeting. One-tailed unpaired t-test (B, G). Two-tailed t-test (C, E, F). Error bars = SEM. (A, D) *N*=1. (B, C, E, F, G) *N*=3. See *also Supplementary Fig. 4 and 5*.

A previous study indicated that wt-p53 regulation of SLC7A11 was context dependent^32^. To this point, we found that p53 knockdown decreased SLC7A11 protein in two CCLs with endogenous wt-p53 expression, HCT116 and U2OS (**Figure 5D**). *SLC7A11* mRNA was decreased following p53 ablation, whilst other NRF2 targets remained unchanged (**Figure S 4C**). This resulted in increased sensitivity to APR-246, however this was response was more pronounced in the HCT116 cells (**Figure 5E**).

Probing deeper into the mechanism underlying wt-p53 regulation of SLC7A11, in contrast to previous studies^30,31^, chromatin-immunoprecipitation (ChIP) studies in our hands showed no enrichment of p53 at the *SLC7A11* promoter (**Figure 5G**). This finding was supported by analysis of 26 publicly available genome-wide p53 chromatin binding profiles that found no p53 occupancy at the *SLC7A11* promoter (www.targetgenereg.org^33^). We also investigated other possible mechanisms of *SLC7A11* suppression relating to the p53 transcriptional network. A comprehensive meta-analysis of the p53 transcriptional program suggested that p53-dependent gene repression relies on the p21-DREAM axis^34^. However, siRNA-mediated ablation of p21 did not restore *SLC7A11* mRNA expression following p53 induction (**Figure 5F**). Together, these results highlight that wt-p53 regulation of *SLC7A11* is indirect, however the suppression of SLC7A11 by altering wt-p53 levels results in increased sensitivity to APR-246.

Finally, analysis of the *SLC7A11* promoter region and previously published c-Myc ChIP-seq data revealed a putative c-Myc binding site and c-Myc occupancy at the *SLC7A11* promoter (**Figure S 5A**). Furthermore, knockdown of c-Myc in HCT116 and U2OS cells resulted in ablation of SLC7A11 protein expression and concomitantly diminished cystine uptake (**Figure S 5B,C**), although this did not decrease the total basal levels of intracellular GSH (**Figure S 5D**). Nevertheless, knockdown of c-Myc resulted in marked increased sensitivity to APR-246 in HCT116 and U2OS (**Figure S 5E**). This suggests that capacity for cystine uptake through SLC7A11 may be the important predictor of APR-246 sensitivity and not necessarily intracellular basal GSH levels *per se*. Together, this highlights that altering SLC7A11 through silencing c-Myc results in increased sensitivity to APR-246 in wt-p53 cancer cell lines, further demonstrating that SLC7A11 expression is a consistent determinant of sensitivity to APR-246 – independent of *TP53* mutation.

## DISCCUSION

Evaluation of APR-246 is currently underway in a phase III clinical trial in *TP53-* mutated MDS in combination with Azacitidine^35^, with preliminary phase II results reporting strong efficacy^36,37^. Whilst previous reports have defined *TP53* mutation status as the presumptive clinical biomarker for APR-246 patient selection^11^, here, we propose that the major predictor for response to APR-246 is expression levels of the cystine-glutamate antiporter, SLC7A11: where low levels correlate with greater sensitivity. Stressing this point further, diminishing the levels of SLC7A11 through altering levels of four key proteins involved in cancer biology, MDM2, ATF4, wt-p53 and c-Myc, increased sensitivity to APR-246 in p53-null/wt-p53 CCLs (**Figure 5H**). Furthermore, whilst loss of ARID1A^8,9^ and mut-p53 protein accumulation^4^ have been shown to suppress SLC7A11 through NRF2 regulation, only *KEAP1* mutation was found to be significantly predictive of resistance to APR-246 across CCLs. As a result, our study critically demonstrates the necessity for broadening the clinical investigation of APR-246 therapy to beyond *TP53* mutant malignancies – in both solid and haematological settings. The clinical impetus for this is highlighted through the results of our clinical investigation into APR-246 in metastatic oesophageal cancer, which indicated that patients without *TP53* mutations may respond to APR-246.

On the point of mut-p53 reactivation by APR-246, notably, Cys277 was recently identified to be a key residue in the DNA binding domain of mut-p53 for MQ docking to drive thermostabilisation^3^. However, here, and in previous studies^4,11^, we have shown that the oesophageal CCL FLO-1, which expresses mut-p53^C277F^, is sensitive to APR-246. In our investigation into the capacity of APR-246 and PX-12 to induce mut-p53 thermostability in cells, APR-246 did not thermostabilise mut-p53 to a greater degree than PX-12. Therefore, the similarity between APR-246 and PX-12 activity in CCLs could be explained by the capacity of these compounds to deplete GSH: PX-12 through thioredoxin inhibition, and APR-246 through direct conjugation via MQ.

Pan-cancer lineage analysis highlighted PNS tumours as a lineage of interest for APR-246, as they express low levels of *SLC7A11* and are highly sensitive to PRIMA-1 (**Figure 2C, 3E**). This is supported by pre-clinical investigations of APR-246 in neuroblastoma models, which demonstrated caspase-independent, p53-independent cell death through glutathione depletion^38^. Given that neuroblastoma is a heterogeneous childhood tumour, with limited therapeutically targetable genomic alterations and poor prognosis, more work investigating the clinical utility of APR-246 in neuroblastoma should be undertaken.

On the strength of our new findings of the value of SLC7A11 as a prognostic marker for APR-246 efficacy, validation is now warranted in patient samples from clinical trials of this drug. Determining SLC7A11 expression in clinical samples by immunohistochemistry^39^, or *in situ* tumour system xc activity by PET imaging using (4S)-4-(3-[^18^F]Fluoropropyl)-Lglutamate (^18^F-FSPG)^40,41^, is achievable and rapidly translatable into the clinic.

## Acknowledgements

This work was supported by a National Health and Medical Research Council (NMHRC) Project Grant (APP1120293) to W.A.P., N.J.C.), a Grand-In-Aid (APP1156945 to N.J.C., W.A.P.) and a Translational Research Project Grant (TRP15012 to W.A.P., N.J.C.). N.J.C. is supported by a Victorian Cancer Agency Fellowship (MCRF16002). K.M.F. is supported by an Australian Research Training Program (RTP) Scholarship. C.S.C is the recipient of, and support by, The Alan & Kate Gibson Research Fellowship, 2018.

## Author Contributions

Conceptualisation: K.M.F., N.J.C.; Methodology: K.M.F., M.C.B., C.S.C, B.Z.Z., H.S.K., D.S.L, K.J.S., N.J.C.; Investigation: K.M.F, M.C.B, C.S.C, B.Z.Z., H.S.K, D.S.L,; Writing – Original draft: K.M.F.; Writing – Review & editing: K.M.F., B.Z.Z., S.H., W.A.P., N.J.C; Resources: K.J.S., Y.H., W.A.P., N.J.C.; Supervision: S.H., W.A.P., N.J.C, Project Administration: N.J.C; Funding acquisition: W.A.P, N.J.C.

## Materials and methods

### Clinical Trial

All 5 patients enrolled in the study received Dose Level 1, which included 75 mg/kg (Lean body mass) APR-246, 750 mg/m^2^/day 5-FU and 25 mg/m^2^ cisplatin. APR-246 was given for 4 days via intravenous infusion over 6 hours starting day 1, 5-FU was administered as a continuous infusion over 96 hr commencing day one, and cisplatin was commenced on day 2 via intravenous infusion over 1 hr following the infusion with APR246. Patient blood was collected before administration of the treatment regimen (Baseline) and between successive cycles of treatment after two cycles. Oesophageal cancer specific ctDNA panel was performed following previously established protocols^13^. Detailed protocols used for the phase Ib/II study can be found here: https://www.clinicaltrials.gov/ct2/show/NCT02999893.

### Compounds and reagents

APR-246 was provided by Aprea Therapeutics. PX-12 was from SelleckChem. Ponasterone-A was from Sigma-Alrich.

### Cell cultures

H1299, HCT116, U2OS and HEK-293T were from ATCC. All cells were maintained at 37°C with 5% CO2. Unless otherwise specified, all culture media contained 10% foetal bovine serum supplemented with 50 U ml^-1^ penicillin and 50 mg ml^-1^ streptomycin (Life Technologies). HCT116, U2OS and HEK-239T cells were grown in DMEM containing 2.5 mM L-glutamine and 4.5 g l^-1^ D-glucose (Life Technologies). H1299 cells were grown in RPMI 1640 medium containing 2.5 mM L-glutamine.

### Boutique siRNA screen

A detailed description of the methods used in the siRNA screen have been previously described (Ref). Briefly, H1299 parental and H1299 overexpressing p53^R273H^ were reverse transfected on day 1, media changed and replaced on day 2 with 10 μM APR-246 or vehicle, and on day 5 cells were fixed and stained with DAPI. 40 nM final concentration of SMARTpool siRNAs (Horizon Discovery) were used and siRNA mixtures. BioTek EL406 washer dispenser and Caliper Sciclone ALH3000 liquid handler were utilised for dispensing the siRNA mixtures and cells. Cells were imaged using the ArrayScan VTI high-content system (ThermoFisher Scientific) and Cellomics Morphology V4 Bioapplication was used to determine cell number based on DAPI staining. Cell numbers were normalised to vehicle + ON-TARGETplus nontargeting control.

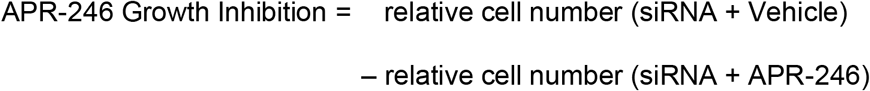

### Cell proliferation and dose-repose assays

For dose-response assays, 10-point log2 serial dilutions of APR-24 and PX-12 were added to 96-well plates containing cells. After 72 h incubation, cell viability was determined using AlamarBlue reagent (Life Technologies) and fluorescence was read at 550 nm/590 nm on a Cytation 3 Imaging Reader (BioTek). For proliferation assays, confluency in 96-well plates was tracked over 72 h using Incuctye FLR (Essen BioScience) following manufacturer’s guidelines.

### Cellular Thermal Shift Assay (CETSA)

Performed as per established protocols^22^. Briefly, in 15 cm plates, H1299 p53^R273H^ cells were plated overnight and dosed for 2 h with pre-heated (90°C for 20 mins in water) 50 μM APR-246 and 25 μM PX-12 or DMSO control. Cells were collected in ice-cold PBS and evenly distributed into strip tubes. Cells were subjected to three freeze-thaw cycles and then the lysates were subjected to indicated temperatures for 3 minutes. Lysates were centrifuged, and supernatants were resolved by SDS-PAGE and analysed western blot (see below).

### Inducible p53 expression

PonA-inducible wt-p53 H1299 cells were a gift from Dr. Paul Neilsen (University of Adelaide). Briefly, H1299 cells were stably transfected with pVgRXR aided by selection in 100 μg ml^-1^ zeocin, followed by stable transfection of pI-TK-Hygro-p53-wt and subsequent selection of clones in 600 μg ml^-1^ hygromycin-B. In all experiments, 5 × 10^5^ cells were seeded in 6 cm plates and wt-p53 expression was induced by 2.5 μg ml^-1^ PonA treatment.

### Genetic knockdown using siRNA

Cells were reverse transfected with 40 nM *SLC7A11, TP53, MYC, CDKN1A* or non-targeting control siRNA pools (siGenome SMARTpool, Dharmacon) using Lipofectamine RNAiMAX solution (Life Technologies) according to the manufacturer’s guidelines. Knockdown efficiency was determined by qPCR and/or western blotting.

### SLC7A11 overexpression

*SLC7A11* was ectopically expressed in H1299 using the GE Dharmacon Precision LentiORF pLOC downstream of the CMV promoter and contains turbo-GFP as a reporter gene. The turbo-RFP gene in place of *SLC7A11* was used as a control. Following transduction, GFP-positive H1299 cells were sorted. The expression of *SLC711* was confirmed by qPCR and western blotting.

### Western blot analysis

As per established protocols^4,11^, cells were lysed at 4°C in RIPA buffer (1 mM EDTA, 1% NP-40, 0.5% sodium deoxychlorate, 0.1% SDS, 50 mM sodium fluoride, 1 mM sodium pyrophosphate in PBS) mixed with protease and phosphatase inhibitors (Roche). Equal amounts of protein were boiled, resolved by SDS-PAGE and transferred to PVDF membranes. Membranes were incubated in blocking buffer (5% skim milk in 0.05% TBS-T) for 1 h at room temperature and probed overnight in primary antibody at 4°C. Blots were washed thrice in 0.05% TBS-T, followed by incubation with peroxidase-conjugated secondary antibody (Dako) for 1 h at room temperature. Protein levels were detected using Amersham ECL Western Blotting Detection reagents (GE Life Sciences) or ECL Plus Western blotting substrate kit (ThermoFisher Scientific). Antibodies are detailed in Supplementary Table 1.

### Quantitative RT-PCR

As per established protocols^4,11^, following RNA extraction with NucleoSpin RNA kit (Macherey-Nagel), reverse transcription reaction with Transcriptor First Strand cDNA Synthesis kit (Roche) and SYBR-green qPCR Lightcycler 480 (Roche), gene expression was normalised to GAPDH and ACTB and determined using the ΔΔC_t_ method. Primer sequences are detailed in Supplementary Table 2.

### Chromatin immunoprecipitation

1.25 × 10^6^ PonA-inducible wt-p53 H1299 cells were seeded in 15 cm plates and wt-p53 was induced for 48 h before proceeding to chromatin immunoprecipitation. Protein-DNA conjugates were cross-linked by the addition of methanol-free 1% formaldehyde (Life Technologies) for 10 min. The cross-linking reaction was quenched by the addition of glycine to a final concentration of 0.05M for 5 min. Cells were collected in ice-cold PBS and sequentially lysed in buffer 1 (50 mM HEPES pH 7.5, 140 mM NaCl, 1 mM EDTA, 10% glycerol, 0.5% NP-40, 0.25% TritonX-100), buffer 2 (10 mM Tris pH 8, 200 mM NaCl, 1 mM EDTA, 0.5 mM EGTA) and buffer 3 (10 mM Tris pH 8, 100 mM NaCl, 1 mM EDTA, 0.5 mM EGTA, 0.1% sodium deoxycholate, 0.5% N-laurolysarcosine) with protease inhibitors to enrich for the nuclear fraction. Lysates were sonicated for 35 min (S220 Focused-ultrasonicator, Covaris), followed by precipitation of cellular debris with 10% TritonX-100. An aliquot of lysate was removed for input sample prior to immunoprecipitation. Protein A-Sepharose beads (Invitrogen) were pre-blocked with BSA and sonicated salmon testes DNA. Immunoprecipitations were performed with p53 antibody-agarose conjugate (Santa Cruz Biotechnology, clone sc-6243) or Protein A-Sepharose only at 4°C overnight. Beads were washed five times with LiCl buffer (50 mM HEPES pH 7.5, 500 mM LiCl, 1 mM EDTA, 50 mM NaCl). Protein-DNA conjugates were eluted by incubation in elution buffer (50 mM Tris pH 8, 10 mM EDTA, 1% SDS) at 65°C for 15 min. Reverse cross-linking was performed by the addition of 0.2 mg ml^-1^ RNaseA (Qiagen) and 0.2 mg ml^-1^ Proteinase K (Promega). Samples were analysed by qPCR with primers specific for the promoters of *SLC7A11* and *CDKN1A*. Primers are detailed in Supplementary Table 3.

### Radioactive cystine uptake assay

Following c-Myc knockdown cells were then washed twice in pre-warmed Cystine-free DMEM (high glucose, no glutamine, no methionine, no cystine) to deplete cystine. At this point, in each well the cystine-free-media was replaced with 300μL uptake buffer containing compound and 0.04μCi of L-[1, 2, 1’, 2’-^14^C]-cystine (PerkinElmer) for 5 minutes at room temperature. Cells were then washed thrice with ice-cold cystine-free-media and lysed in 400μL 0.2M NaOH with 1% SDS. This lysate was added to 4mL of scintillation fluid, and radioactive counts per minute were obtained using a scintillation counter. To control for differences in the absolute counts of radioactivity between replicates, data were normalised to DMSO control within each replicate.

### Intracellular glutathione assay

Total intracellular GSH level was assayed using the Cayman Chemicals Glutathione kit as per the manufacturer’s instructions. GSH concentration was calculated from an internal standard curve and normalised to total cell number as determined from parallel plates.

### Statistical analysis

Data were presented as mean ± SEM and analysed by Student’s *t*-test unless otherwise indicated. GI50 dose of single agents was determined by fitting the Hill equation. Statistical analyses and data presentation were performed using Prism 8 (GraphPad). For correlation analysis, the compound activity profiles and *SLC7A11* expression data were obtained from Cancer Dependency Map (https://www.depmap.org) to determine potential associations. Pearson correlations between compound AUC and gene expression were calculated using *cor.test* function in R (Version 3.6.0) for ~8000 cancer cell lines and 481 compounds.

## SUPPLEMENTAL INFORMATION

### SUPPLEMENTAL FIGURE LEGENDS

**Figure S1.**
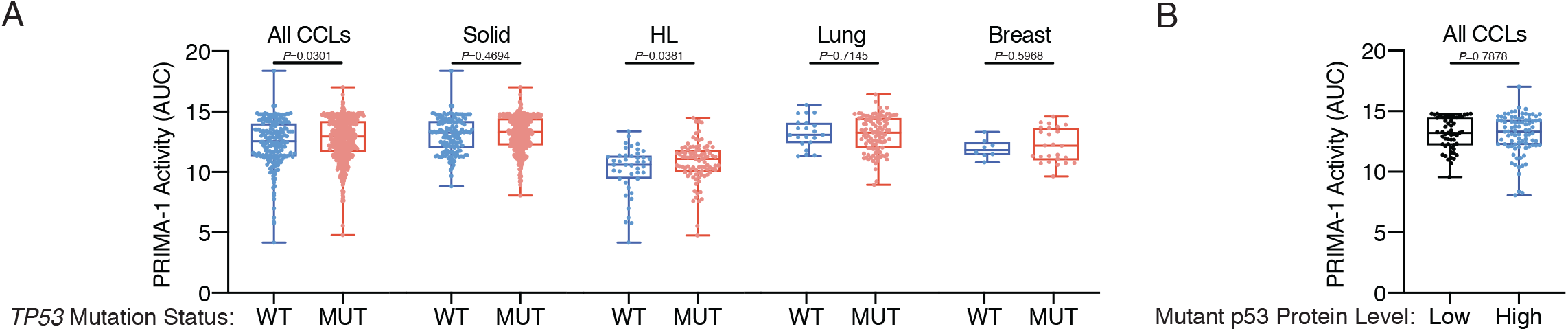
*TP53* mutation alone is not predictive of cancer cell response to PRIMA-1. (A) Boxplots (min-max) of PRIMA-1 AUC across the cancer therapeutics response portal v2 (CTRPv2) stratified by *TP53* mutation status in all cancer cell lines (CCLs), solid, haemopoietic and lymphoid (HL), lung and breast cancer cell lines. (B) Boxplot of PRIMA-1 AUC in all *TP53* mutant CCLs comparing low (z<-1) and high (z>1) mutant-p53 protein levels based on proteomics. Two-way unpaired t-test (A, B).

**Figure S2.**
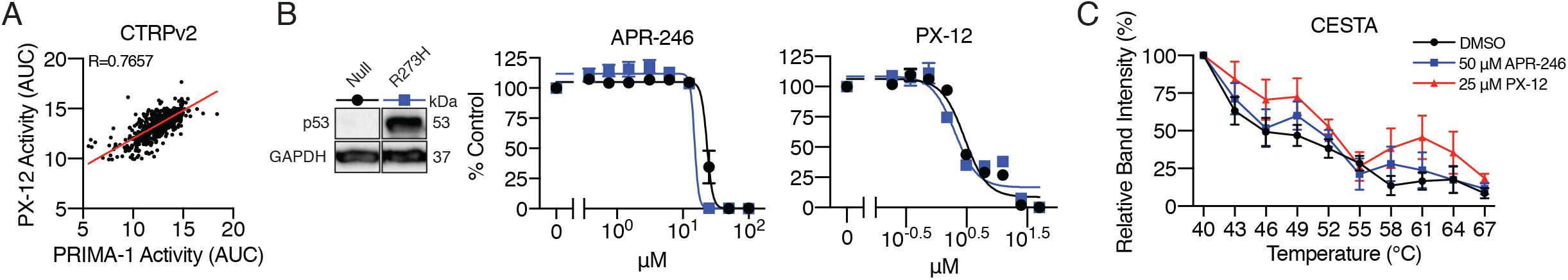
Correlation of APR-246 and PX-12 activity in CTRPv2 and their impact on mutant-p53 thermostability. (A) Scatterplot correlating PRIMA-1 and PX-12 AUC across all CCLs in CTRPv2. (B) Confirmation of mutant-p53 overexpression in H1299 cells (left). Viability following 72 h treatment with APR-246 or PX-12 in p53^Null^ and p53^R273H^ H1299 cells (right). (C) Quantification of cellular thermal stability assay from Figure 2I. Pearson’s correlation (A). Error bars = SEM. (B – western blot) *N*=1. (B – dose responses, C) *N*=3.

**Figure S3.**
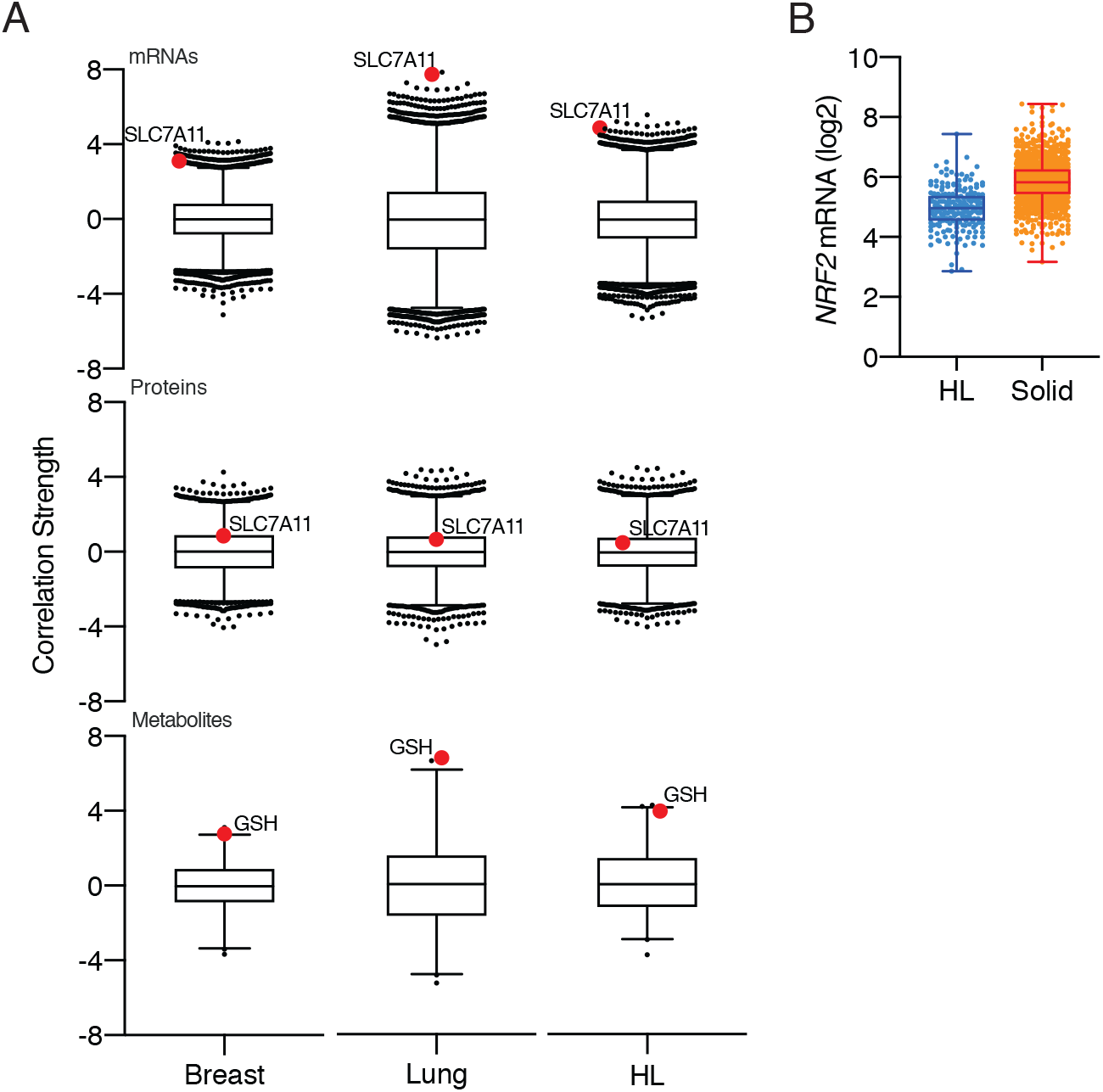
Extended correlation analysis in haematological and lymphoid, breast and lung cancer cell lines. (A) Box-and-whisker plots (1^st^-99^th^ percentile) of Fischer’s transformed z-scored Pearson correlation strengths of PRIMA-1 AUC and gene transcripts, proteins and metabolites in breast or lung cell lines as indicated. Red dots highlight SLC7A11 mRNA and protein, and glutathione (GSH) correlation strengths. (B) Boxplots (min-max) comparing NRF2 mRNA and protein levels in HL and solid tumour derived cell lines.

**Figure S4.**
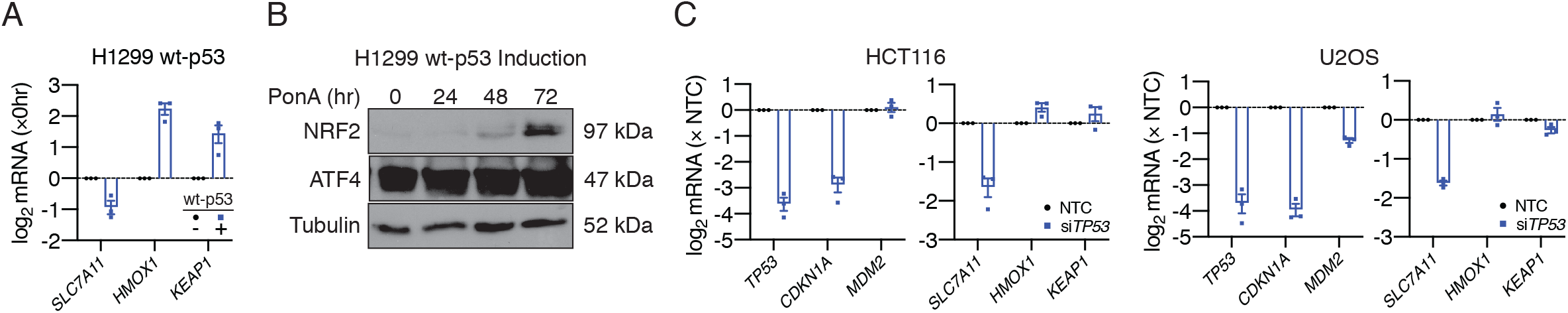
The effects of p53 overexpression and repression on *SLC7A11* mRNA expression. (A) *SLC7A11, HMOX1* and *KEAP1* mRNA levels in H1299 wt-p53 following 48 h of PonA treatment. (B) NRF2 and ATF4 protein levels following induction of wt-p53 with PonA over indicated timepoints. (C) mRNA expression of *TP53*, p53 target genes (*CDKN1A/p21, MDM2*) and NRF2 target genes (*SLC7A11, HMOX1, KEAP1*) following p53 knockdown compared to non-targeting control (NTC) in HCT116 (left) and U2OS (right) cells. Error bars = SEM. *N*=3 (A, C). *N*=1 (B).

**Figure S5.**
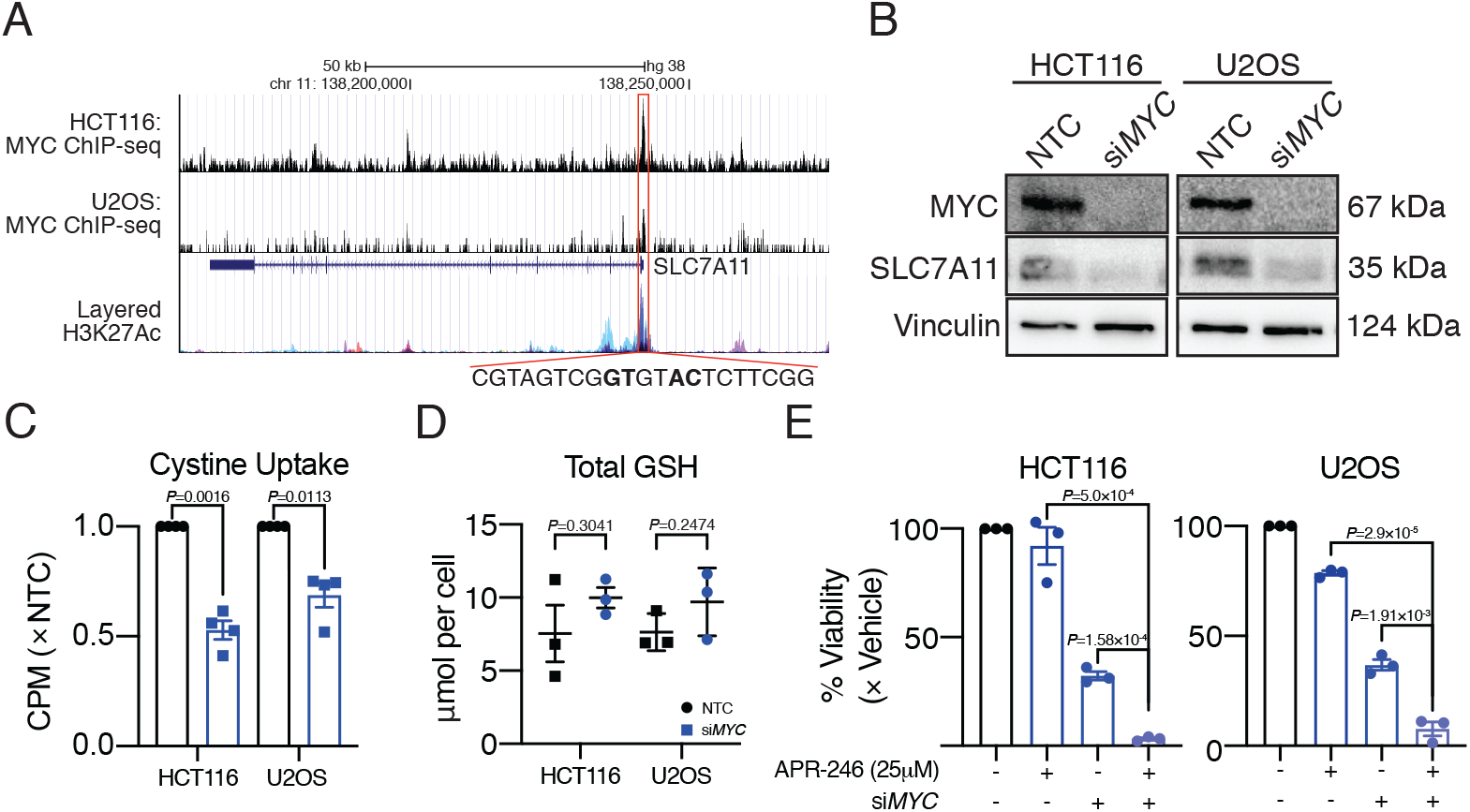
MYC directly regulates SLC7A11 levels and APR-246 sensitivity. (A) Association of MYC with *SLC7A11* promoter in HCT116 and U2OS shown according to the ENCODE ChIP-seq data overlayed with H3K27Ac. Indicates the predicted MYC consensus binding site based on Eukaryotic Promoter Database (EPD) in bold. (B) MYC and SLC7A11 protein, (C) cystine uptake and (D) total GSH levels following *MYC* knockdown in HCT116 and U2OS cells compared to NTC. (E) Viability following 24 h *MYC* knockdown and subsequent treatment with 25 μM APR-246 or vehicle for 72 h. Two-way unpaired t-test (C, D, E). Error bars = SEM. *N=1* (B). *N*=3 (D, E). *N*=4 (C).

## References

1 Bykov, V. J. N., Eriksson, S. E., Bianchi, J. & Wiman, K. G. Targeting mutant p53 for efficient cancer therapy. Nat Rev Cancer 18, 89–102, doi:10.1038/nrc.2017.109 (2018).

2 Lambert, J. M. et al. PRIMA-1 reactivates mutant p53 by covalent binding to the core domain. Cancer Cell 15, 376–388, doi:10.1016/j.ccr.2009.03.003 (2009).

3 Zhang, Q., Bykov, V. J. N., Wiman, K. G. & Zawacka-Pankau, J. APR-246 reactivates mutant p53 by targeting cysteines 124 and 277. Cell Death Dis 9, 439, doi:10.1038/s41419-018-0463-7 (2018).

4 Liu, D. S. et al. Inhibiting the system xC(-)/glutathione axis selectively targets cancers with mutant-p53 accumulation. Nat Commun 8, 14844, doi:10.1038/ncomms14844 (2017).

5 Haffo, L. et al. Inhibition of the glutaredoxin and thioredoxin systems and ribonucleotide reductase by mutant p53-targeting compound APR-246. Sci Rep 8, 12671, doi:10.1038/s41598-018-31048-7 (2018).

6 Krayem, M. et al. p53 Reactivation by PRIMA-1(Met) (APR-246) sensitises (V6OOE/K)BRAF melanoma to vemurafenib. Eur J Cancer 55, 98–110, doi:10.1016/j.ejca.2015.12.002 (2016).

7 Bao, W. et al. PRIMA-1Met/APR-246 induces wild-type p53-dependent suppression of malignant melanoma tumor growth in 3D culture and in vivo. Cell Cycle 10, 301–307, doi:10.4161/cc.10.2.14538 (2011).

8 Ogiwara, H. et al. Targeting the Vulnerability of Glutathione Metabolism in ARID1A-Deficient Cancers. Cancer Cell 35, 177–190 e178, doi:10.1016/j.ccell.2018.12.009 (2019).

9 Sasaki, M. et al. Efficacy of glutathione inhibitors for the treatment of ARID1A-deficient diffuse-type gastric cancers. Biochem Biophys Res Commun 522, 342–347, doi:10.1016/j.bbrc.2019.11.078 (2020).

10 Viswanathan, V. S. et al. Dependency of a therapy-resistant state of cancer cells on a lipid peroxidase pathway. Nature 547, 453–457, doi:10.1038/nature23007 (2017).

11 Liu, D. S. et al. APR-246 potently inhibits tumour growth and overcomes chemoresistance in preclinical models of oesophageal adenocarcinoma. Gut 64, 1506–1516, doi:10.1136/gutjnl-2015-309770 (2015).

12 A Study of APR-246 in Oesophageal Cancer (APROC), <https://www.clinicaltrials.gov/ct2/show/NCT02999893> (2016).

13 Cabalag, C. S. et al. Circulating tumour DNA in oesophageal cancer – utility in tumour staging, monitoring of treatment response and the detection of recurrent disease. Gut (Under Review) (2020).

14 Seashore-Ludlow, B. et al. Harnessing Connectivity in a Large-Scale Small-Molecule Sensitivity Dataset. Cancer Discov 5, 1210–1223, doi:10.1158/2159-8290.CD-15-0235 (2015).

15 Rees, M. G. et al. Correlating chemical sensitivity and basal gene expression reveals mechanism of action. Nat Chem Biol 12, 109–116, doi:10.1038/nchembio.1986 (2016).

16 Basu, A. et al. An interactive resource to identify cancer genetic and lineage dependencies targeted by small molecules. Cell 154, 1151–1161, doi:10.1016/j.cell.2013.08.003 (2013).

17 Itoh, K. et al. Keap1 represses nuclear activation of antioxidant responsive elements by Nrf2 through binding to the amino-terminal Neh2 domain. Genes Dev 13, 76–86, doi:10.1101/gad.13.1.76 (1999).

18 DeNicola, G. M. et al. Oncogene-induced Nrf2 transcription promotes ROS detoxification and tumorigenesis. Nature 475, 106–109, doi:10.1038/nature10189 (2011).

19 Donehower, L. A. et al. Integrated Analysis of TP53 Gene and Pathway Alterations in The Cancer Genome Atlas. Cell Rep 28, 3010, doi:10.1016/j.celrep.2019.08.061 (2019).

20 Nusinow, D. P. et al. Quantitative Proteomics of the Cancer Cell Line Encyclopedia. Cell 180, 387–402 e316, doi:10.1016/j.cell.2019.12.023 (2020).

21 Jordan, B. F. et al. The thioredoxin-1 inhibitor 1-methylpropyl 2-imidazolyl disulfide (PX-12) decreases vascular permeability in tumor xenografts monitored by dynamic contrast enhanced magnetic resonance imaging. Clin Cancer Res 11, 529–536 (2005).

22 Martinez Molina, D. et al. Monitoring drug target engagement in cells and tissues using the cellular thermal shift assay. Science 341, 84–87, doi:10.1126/science.1233606 (2013).

23 Ghandi, M. et al. Next-generation characterization of the Cancer Cell Line Encyclopedia. Nature 569, 503–508, doi:10.1038/s41586-019-1186-3 (2019).

24 Li, H. et al. The landscape of cancer cell line metabolism. Nat Med 25, 850–860, doi:10.1038/s41591-019-0404-8 (2019).

25 Sasaki, H. et al. Electrophile response element-mediated induction of the cystine/glutamate exchange transporter gene expression. J Biol Chem 277, 44765–44771, doi:10.1074/jbc.M208704200 (2002).

26 Bassi, M. T. et al. Identification and characterisation of human xCT that co-expresses, with 4F2 heavy chain, the amino acid transport activity system xc. Pflugers Arch 442, 286–296, doi:10.1007/s004240100537 (2001).

27 Lu, S. C. Regulation of glutathione synthesis. Mol Aspects Med 30, 42–59, doi:10.1016/j.mam.2008.05.005 (2009).

28 Haupt, Y., Maya, R., Kazaz, A. & Oren, M. Mdm2 promotes the rapid degradation of p53. Nature 387, 296–299, doi:10.1038/387296a0 (1997).

29 Riscal, R. et al. Chromatin-Bound MDM2 Regulates Serine Metabolism and Redox Homeostasis Independently of p53. Mol Cell 62, 890–902, doi:10.1016/j.molcel.2016.04.033 (2016).

30 Jiang, L. et al. Ferroptosis as a p53-mediated activity during tumour suppression. Nature 520, 57–62, doi:10.1038/nature14344 (2015).

31 Wang, S. J. et al. Acetylation Is Crucial for p53-Mediated Ferroptosis and Tumor Suppression. Cell Rep 17, 366–373, doi:10.1016/j.celrep.2016.09.022 (2016).

32 Xie, Y. et al. The Tumor Suppressor p53 Limits Ferroptosis by Blocking DPP4 Activity. Cell Rep 20, 1692–1704, doi:10.1016/j.celrep.2017.07.055 (2017).

33 Fischer, M., Grossmann, P., Padi, M. & DeCaprio, J. A. Integration of TP53, DREAM, MMB-FOXM1 and RB-E2F target gene analyses identifies cell cycle gene regulatory networks. Nucleic Acids Res 44, 6070–6086, doi:10.1093/nar/gkw523 (2016).

34 Fischer, M. Census and evaluation of p53 target genes. Oncogene 36, 3943–3956, doi:10.1038/onc.2016.502 (2017).

35 APR-246 & Azacitidine for the Treatment of TP53 Mutant Myelodysplastic Syndromes (MDS), <https://clinicaltrials.gov/ct2/show/NCT03745716> (2018).

36 Sallman, D. A. et al. Phase 2 Results of APR-246 and Azacitidine (AZA) in Patients with TP53 mutant Myelodysplastic Syndromes (MDS) and Oligoblastic Acute Myeloid Leukemia (AML). Blood 134, 676–676, doi:10.1182/blood-2019-131055 (2019).

37 Cluzeau, T. et al. APR-246 Combined with Azacitidine (AZA) in TP53 Mutated Myelodysplastic Syndrome (MDS) and Acute Myeloid Leukemia (AML). a Phase 2 Study By the Groupe Francophone Des Myélodysplasies (GFM). Blood 134, 677–677, doi:10.1182/blood-2019-125579 (2019).

38 Mlakar, V. et al. PRIMA-1(MET)-induced neuroblastoma cell death is modulated by p53 and mycn through glutathione level. J Exp Clin Cancer Res 38, 69, doi:10.1186/s13046-019-1066-6 (2019).

39 Wang, W. et al. CD8(+) T cells regulate tumour ferroptosis during cancer immunotherapy. Nature 569, 270–274, doi:10.1038/s41586-019-1170-y (2019).

40 Mittra, E. S. et al. Pilot Preclinical and Clinical Evaluation of (4S)-4-(3-[18F]Fluoropropyl)-L-Glutamate (18F-FSPG) for PET/CT Imaging of Intracranial Malignancies. PLoS One 11, e0148628, doi:10.1371/journal.pone.0148628 (2016).

41 Park, S. Y. et al. Clinical Evaluation of (4S)-4-(3-[(18)F]Fluoropropyl)-L-glutamate ((18)F-FSPG) for PET/CT Imaging in Patients with Newly Diagnosed and Recurrent Prostate Cancer. Clin Cancer Res 26, 5380–5387, doi:10.1158/1078-0432.CCR-20-0644 (2020).

